# Exploring Neural Dynamics in the Auditory Telencephalon of Crows using Functional Ultrasound Imaging

**DOI:** 10.1101/2024.11.26.625563

**Authors:** Diana A. Liao, Eva Schwarzbach, Andreas Nieder

## Abstract

Crows, known for advanced cognitive abilities and vocal communication, rely on intricate auditory systems. While the neuroanatomy of corvid auditory pathways is partially explored, the underlying neurophysiological mechanisms are largely unknown. This study used functional ultrasound imaging (fUSi) to investigate sound-induced cerebral blood volume (CBV) changes in the field L complex of the crow’s auditory telencephalon, a functional analogue to the mammalian auditory cortex. FUSi revealed frequency-specific CBV responses, showing a tonotopic organization within the field L complex, with low frequencies in the posterior-dorsal region and high frequencies in the anterior-ventral region, similar to neuronal patterns reported in other songbirds. The responses were more robust when the crows were awake compared to when they were anesthetized. Machine learning analyses showed fUSi signals could be used to classify sound types accurately. Shorter stimuli reliably triggered transient CBV increases, while longer sounds resulted in variable responses, including negative deflections. This variability in CBV responses suggests a delineation of subregions within the field L complex with the central region (L2) as the primary processing hub and outer regions (L1, L3) integrating auditory information. These findings highlight the potential of fUSi for providing high-resolution insights into functional systems in corvids, enabling future exploration of task-related cognitive dynamics.

## Introduction

Corvids—crows, ravens, and jays—are the largest members of the songbirds and are known for their exceptional cognitive abilities (Emery and Clayton, 2004; Taylor, 2014; Nieder, 2023). Corvids utilize their behavioral flexibility to engage in sophisticated audio-vocal communication. They possess an impressive ability to produce a diverse range of sounds, including the imitation of human speech sounds (Coombs, 1960; Chamberlain and Cornwell, 1971; Brown, 1985; Webber and Stefani, 1990; Bluff et al., 2010). Corvids strategically use their vocalizations to navigate complex social dynamics, signaling identity, familiarity, dominance, and group membership, thus demonstrating sophisticated acoustic social recognition (Hopp et al., 2001; Kondo et al., 2010; Wascher et al., 2012; Mates et al., 2015; Szipl et al., 2017; Cunha and Griesser, 2021; Martin et al., 2024). In particular, crows exhibit elaborate cognitive control over their vocalizations, a skill supported by precise audio-vocal feedback mechanisms (Brecht et al., 2019; 2023; Liao et al., 2024). These remarkable traits are underpinned by corvids’ advanced auditory processing capabilities.

The corvids’ auditory system, known in birds as the field L complex—analogous to the mammalian auditory cortex—plays a key role in processing and interpreting sounds. Recent research has begun to uncover the neuroanatomy of the telencephalic audio-vocal pathways in crows (Kersten et al., 2021; 2022; 2024; Moll et al., 2024), but the neurophysiological mechanisms remain unexplored. Additionally, communicative behaviors emerge from dynamic interactions of distributed brain areas such as those for vocal perception (e.g. the field L complex) and those for vocal production (e.g. the song system). Thus, a neuroimaging method with a large field of view is critical to help uncover the substrates for vocal communication. Here, we aim to investigate the auditory telencephalon of crows using neuroimaging, specifically functional ultrasound imaging (fUSi), an emerging technique not yet applied to songbirds.

Neuroimaging in humans primarily relies on functional magnetic resonance imaging (fMRI). The main advantages of fMRI are its ability to non-invasively capture high-resolution, real-time, and brain-wide activity by measuring changes in blood oxygenation. In recent years, fMRI has also been successfully applied to birds, including finches (Van Ruijssevelt et al., 2013; 2018), pigeons (Behroozi et al., 2020; Ungurean et al., 2023), and chickens (Behroozi et al., 2024). However, the technical complexity, spatio-temporal resolution, and the need for subjects to remain completely still during measurements still pose significant challenges for neuroimaging in behaving birds and other small animals. To date, fMRI in awake birds performing controlled and complex tasks necessary to study cognitive functions or vocal production has yet to be achieved.

Functional ultrasound imaging (fUSi) is an emerging neuroimaging technique that overcomes some limitations of fMRI by using ultra-fast, high-frequency sound waves to measure brain activity through detailed images of blood volume dynamics (Macé et al., 2011, Urban et al., 2015). Offering high spatial resolution (∼100 µm) and temporal resolution in the hundreds of milliseconds, fUSi uses lightweight probes to map brain activity by emitting and recording sound waves that reflect off moving red blood cells. The small, head-mounted probes make it especially well-suited for use in awake, behaving animals during behavioral tasks (Urban et al., 2015, Sieu et al., 2015, Tiran et al., 2017, Takahashi et al., bioRxiv, El Hady et al., 2024). Since fUSi does not rely on magnetic fields, unlike fMRI, it could be more easily combined with other neurophysiological methods, like electrophysiology (Nunez-Elizalde et al., 2022, Lambert et al., bioRxiv, Claron et al., 2023), calcium imaging (Aydin et al., 2020), pharmacological manipulations (Di Ianni et al., 2024), or optogenetics (Edelman et al., 2021).

To date, fUSi has been employed in only one bird species, the pigeon (Rau et al., 2018). This proof-of-concept study demonstrated that fUSi enables the investigation of brain responses to both visual and auditory stimulation. In the current study, we leveraged fUSi to assess its suitability and sensitivity in a corvid songbird, the carrion crow (*Corvus corone)*. Specifically, we explored responses to different acoustic stimuli in the corvid field L complex, a key auditory thalami-recipient input and telencephalic processing region in birds.

## Results

We positioned the ultrasound probe (IcoPrime-15 MHz, Iconeus, Paris, France) along an anterior-posterior axis above the partial craniotomy window; the region covered includes the auditory telencephalon termed the field L complex (**Fig. 1A, B**) The partial craniotomy covered most of the posterior left hemisphere, crossing slightly over the midline into the right hemisphere. The probe imaged a 2-dimensional (2D) brain slice with dimensions of 14 mm anterior-posterior by 19 mm dorso-ventral (**Fig. 1C, D**).

**Figure 1:**
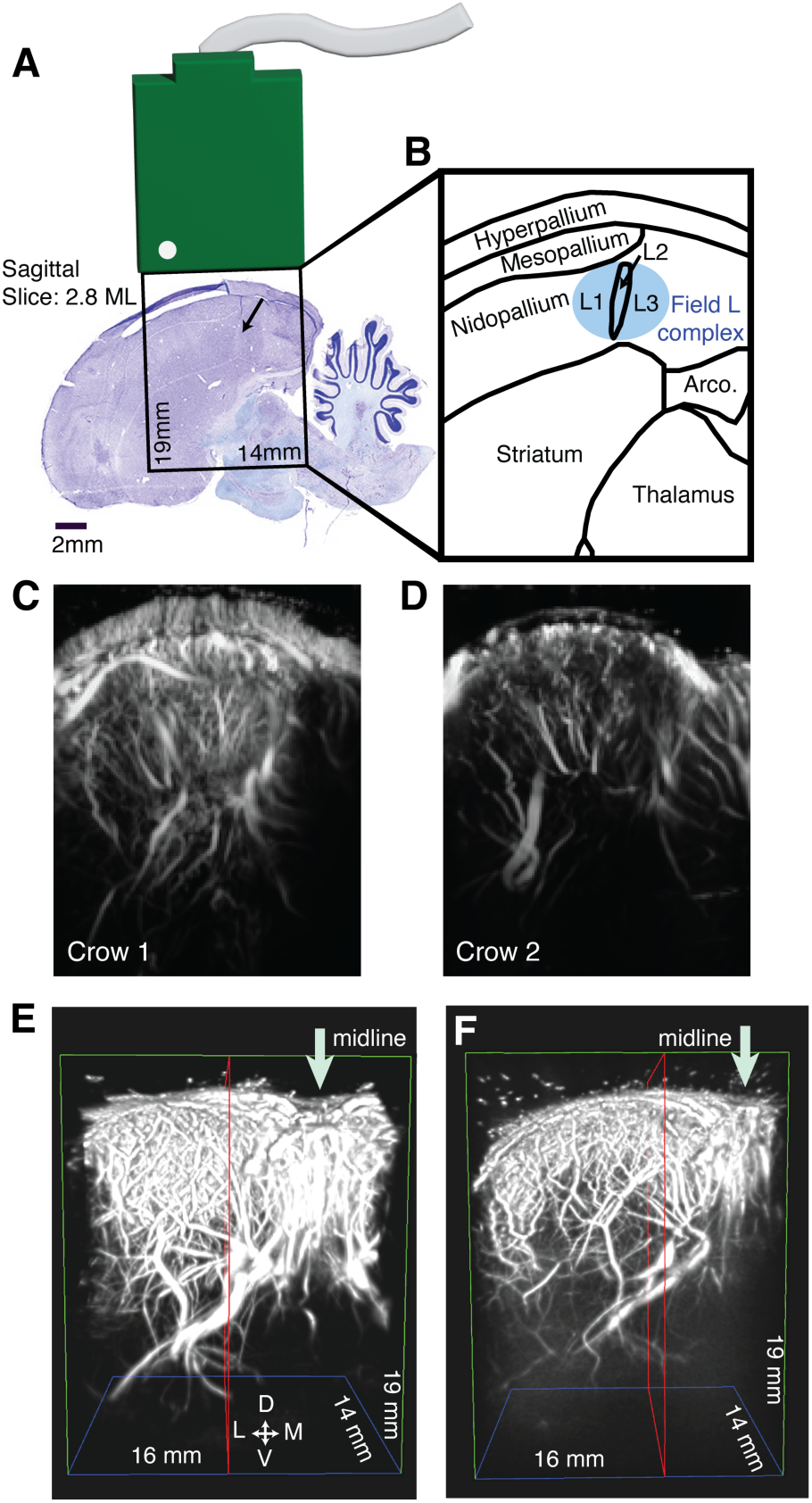
Ultrasound imaging of the posterior telencephalon in the crow. **A)** Sagittal histological section of a crow brain (adapted from the Crow Brain Atlas by Kersten et al., 2022), showing the ultrasound probe positioned over the posterior half of the telencephalon (with the left side representing the anterior direction, 2.8 mm lateral from the midline). The black outline represents the imaging window (14 mm x 19 mm) extending into the brain. **B)** Magnified schematic of the anatomical structures found within the imaging window depicted in A. The avian auditory telencephalon, specifically the field L complex is shaded in blue. L2, a subregion of the field L complex, is highlighted based on its fiber characteristics (marked by arrow in **A**). **C)** A 2D angiogram (visualization of blood vessels) for crow 1, acquired from the imaging window depicted at the coordinates outlined in **A** and **B**. **D)** 2D angiogram for crow 2. **E)** A 3D angiogram of the left hemispheric posterior telencephalon of crow 1, mildly extending across the midline (indicated by a downward-pointing arrow) into the left hemisphere. This image displays the brain volume beneath the partial craniotomy window, generated by graphically combining 2D images (one scan every 200 µm in medio-lateral extension), as shown in **C** and **D**. **F)** 3D angiogram of crow 2.

The first step was to image the brain structures beneath the partial craniotomy. Using a linear motor stage (SLC – 1740, SmarAct GmbH motor) to precisely move the ultrasound probe from medial to lateral in 200 µm steps, we acquired a 2D image at each position. The high spatial resolution of ultrasound imaging enabled detailed depiction of even fine blood vessels branching from the major vessels, which predominantly run along the dorso-ventral axis (**Fig. 1C, D**). These individual 2D images were graphically combined to create a 3-dimensional (3D) image of the brain volume (**Fig. 1E, F**) that reveals the fine details of the brain vasculature network.

### Functional localizer protocol

To test for auditory-related changes in cerebral blood volume (CBV) to localize the putative avian auditory telencephalon (i.e. the field L complex), we used an auditory localizer protocol applied to anesthetized crows. The ultrasound probe was moved in the medio-lateral direction in 1 mm steps within the area of the partial craniotomy. At each position, auditory noise bursts (5 second duration, surrounded by 5 seconds of silence) were presented to test for stimulus-correlated changes in CBV. Online, using dedicated software (IcoStudio), CBV responses for each voxel were correlated with the stimulus vector (noise sound) to create correlation maps (**Fig. 2A**). We visualized correlated CBV responses by thresholding activity to 20% to see which slices contained the most correlated responses.

**Figure 2:**
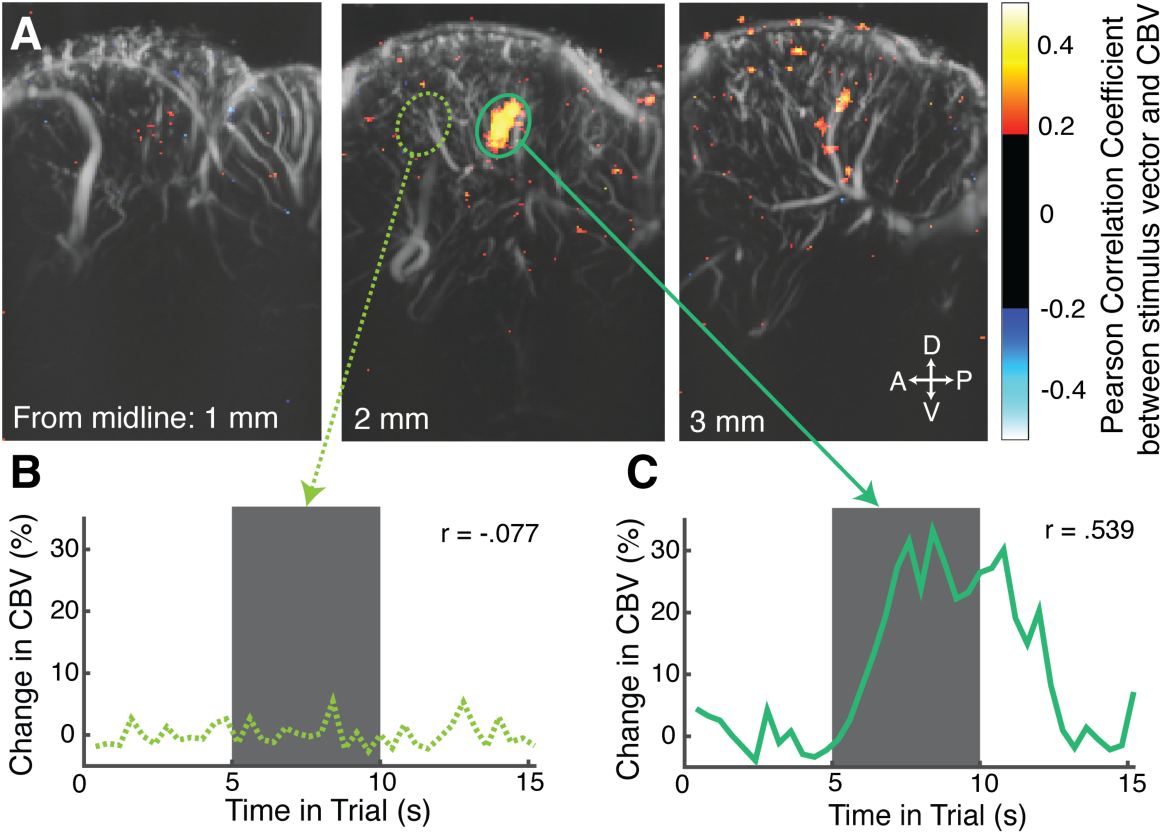
Localizer protocol for auditory activity in anesthetized crows. **A)** Functional ultrasound imaging was performed while stimulating with noise bursts and moving the probe from medial to lateral. Brain slices from crow 2 at 1 mm, 2 mm, and 3 mm lateral to the midline are shown as examples. The middle slice, at 2 mm lateral from the midline, shows a pronounced correlation of the CBV signal with the noise stimulus in a brain region that coincides with the field L complex depicted in **Fig. 1B. B)** Example CBV trace (in % relative to the silence baseline) during a noise burst in a region at slice 2 mm lateral from the midline that showed no stimulus-correlated change. This activity trace is taken from the brain region in **A** (middle) indicated by a dotted light green circle. The gray area marks the noise stimulation period. **C)** Example CBV trace during a noise burst at the same slice that is stimulus correlated. This activity trace is taken from voxels in the brain region indicated by a solid green circle.

In crow 2, a slice 2 mm lateral from the midline showed a pronounced activated region at a location that coincided with the core of the field L complex, as predicted by neuroanatomical coordinates (Kersten et al., 2022) (**Fig. 2A**, middle panel). Within this region of activation, the CBV increased sharply about 2 seconds after the physical sound onset, reached a plateau of roughly 30% signal increase (100% signal change indicates that CBV has doubled compared to baseline), and returned to baseline with a delay comparable to the onset delay (**Fig. 2C**). In a region outside of this ‘hot spot,’ but neighboring it, the CBV remained at baseline throughout sound stimulation (**Fig. 2B**). Slices positioned more medially or laterally showed less stimulus-correlated CBV activation. The slice with the most pronounced activation during the functional localizer protocol (2 mm lateral from the midline in crow 2, and 3 mm lateral from the midline in crow 1) was used to secure the probe at that particular location to be subsequently used for the following stimulation protocols, measurements, and analyses.

### Responses to bird vocalizations

To test whether the CBV changes were specific and robust enough to represent complex sounds, we employed a block design with playbacks of three different acoustically rich vocalizations from a crow, a pigeon, and a canary. The vocalizations were intensity-equalized but naturally differed in other acoustic dimensions, such as frequency range (**Fig. 3A**, pigeon: low frequencies; crow: middle frequencies; canary: high frequencies). Each call sequence consisted of 5-second intervals of each repeated vocalization, with 15 seconds of silence between each vocalization block.

**Figure 3:**
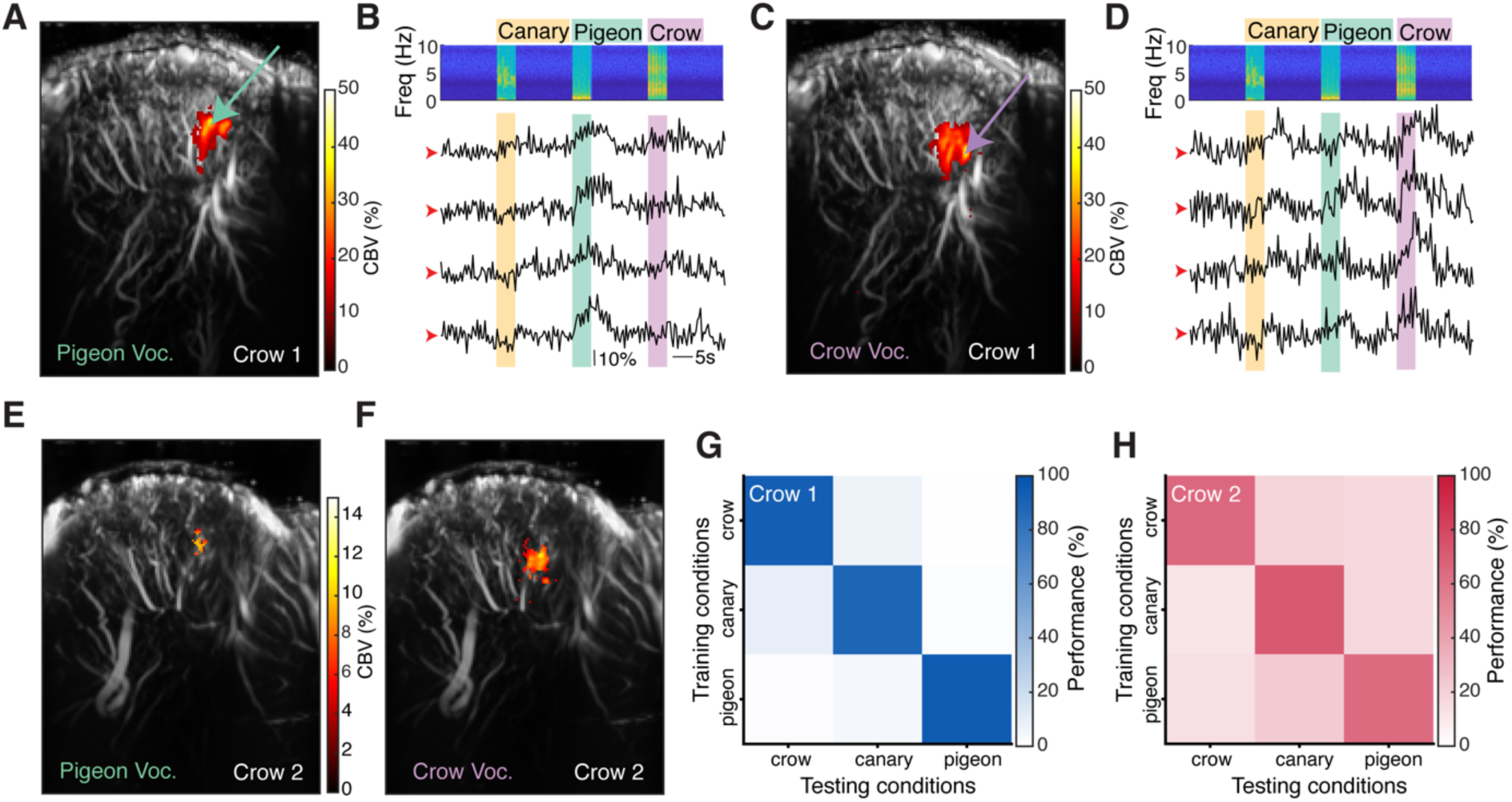
Mapping CBV changes to playback of complex bird vocalizations. **A)** Bird vocalization stimulation sequence and corresponding CBV changes from an example voxel responsive to pigeon sounds in crow 1. The top panel shows the spectrogram of one sequence of bird vocalizations (canary, pigeon, crow). The four bottom panels depict the single-trial CBV responses from four repetitions of this sequence. The CBV traces were taken from a voxel indicated by the green arrow in **B**. **B)** Mean activation map in crow 1 for pigeon vocalizations (green stimulus interval shown in **A**). **C)** Vocalization sequence and corresponding CBV changes from a voxel responsive to crow sounds. The CBV traces were taken from a voxel indicated by the purple arrow in **D**. **D)** Activation map for crow vocalizations (purple stimulus interval shown in **C**). **E)** Activation map for pigeon vocalizations in crow 2. **F)** Activation map for crow vocalizations in crow 2. **G)** Confusion matrices showing the performance of an MVPA classifier predicting the type of bird vocalization played based on voxel activity for crow 1 (in blue). **H)** Confusion matrices showing the performance of an MVPA classifier for crow 2 (in red).

**Figure 3A** shows four repetitions of one vocalization sequence and the temporally correlated CBV traces obtained from an example voxel indicated in Figure 3B. The CBV changes align with the stimulation onsets, with the largest CBV responses occurring during pigeon vocalizations. CBV responses for each voxel were correlated with the hemodynamic response convolved stimulus vector (p<.001, FDR corrected) for each bird vocalization select active voxels in order to generate the mean activity map. The pigeon map is shown in Figure 3B where the resulting activation was located primarily in a dorsal region.

Figure 3C shows four repetitions of the same vocalization sequence as above and the temporally correlated CBV traces obtained from an example voxel indicated in Figure 3D, which was most responsive to crow vocalizations. The activation map for crow vocalizations (Fig. 3D) was located more ventrally compared to the map for pigeon vocalizations (Fig. 3B). Similar regionally different activation patterns were seen in crow 2 to pigeon vocalizations (Fig. 2E) and crow vocalizations (Fig. 2F).

Next, we applied machine-learning classifier analyses to the fUSi data. Specifically, we performed a multivoxel pattern analysis (MVPA) to infer the type of auditory responses in the imaged brain area. MVPA is a supervised classification technique used to identify relationships between spatial-temporal patterns of fUSi activity and stimuli conditions. Figure 3G shows the classifier performance (plotted as a confusion matrix) in predicting vocalizations based on data recorded from crow 1. The accuracy, i.e. the proportion with which the classifier correctly predicted a vocalization condition, is shown along the main diagonal of the confusion matrix. The classifier accuracy for crow 1 is at 91.9 ± 3.67% and thus significantly above the chance level of 33% correct classification, given the 3 possible vocalization conditions (one-sample t-test, p < 0.001, t(19) = 20.488). For crow 2 the classifier was also able to predict all three vocalizations with above chance accuracy (mean: 66.7 ± 3.84%; one-sample t-test, p < 0.001, t(19) = 13.036, Fig. 3H).

These data demonstrate that the CBV changes measured with fUSi were specific and robust enough to represent complex sounds. Moreover, the distinct activation locations of the bird vocalizations along the dorso-ventral axis suggest that different sound frequencies within the vocalizations activate specific regions of the auditory telencephalon. Based on this, we next investigated the tonotopic mapping within the field L complex in greater detail.

### Tonotopic mapping in awake crows using pure tones

For the investigation of tonotopic mapping, we presented the crows with six pure tone sounds of equal intensity, with frequencies increasing in 1-octave steps (250 Hz, 500 Hz, 1000 Hz, 2000 Hz, 4000 Hz, and 8000 Hz). These frequencies are known to cover the typical hearing range of crows and other songbirds (Dooling et al., 2000; Jensen and Klokker, 2006). The six sounds were presented for 5 seconds each in a pseudo-randomized order in each scan (Fig. 4A). After each sound, 15 seconds of silence followed. Figure 4A shows example CBV traces averaged over nine voxels in the field L complex. The responses to specific sound frequencies (indicated by color bars) were similar between scans. For further analysis, the traces were aligned at the onset of the stimulation period, with the five seconds before stimulation onset taken as the individual baseline.

**Figure 4:**
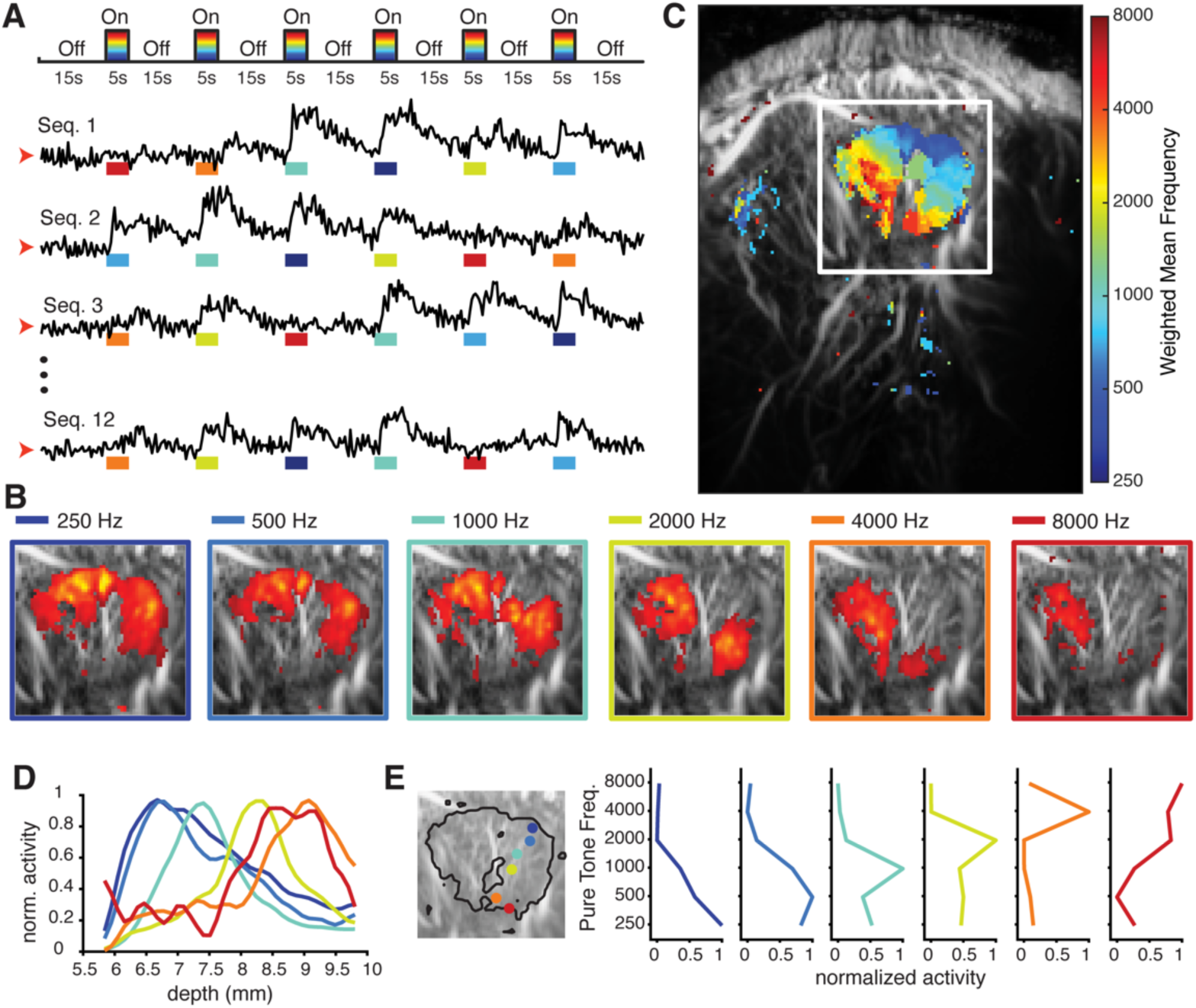
Topographic arrangement of pure tone representations in awake crows. **A)** The top panel schematic shows the pure tone sequence, consisting of 6 sequential pure tones (5s duration) of systematically varying frequencies (color coded) separated by silence (15s duration). The CBV traces below are from voxels in the field L complex during four example scans temporally aligned to the sound sequence. **B)** Average activation maps in crow 1 (graphically zoomed in on the field L complex; see white square in **C**) arranged in increasing sound frequencies from left to right. The color of the frames indicates the sound frequency. **C)** Tonotopic frequency map: Activation to each sound frequency was superimposed and used to compute a weighted average, with cool to warm colors representing increasing sound frequencies. **D)** Iso-intensity frequency tuning curves of the voxels along the dorso-ventral imaging depth axis. CBV activity was normalized to each frequency’s maximum response. **E)** Zoomed-in square of the field L complex (significant voxel cluster outlined in black) with example voxels superimposed on image. Next to image, the 6 example voxels are shown with tuning for a particular frequency. Normalized activity is on the X axis and frequency is shown on the Y axis.

We calculated the normalized activity of each voxel for each sound frequency presentation. The resulting change in CBV is relative to the baseline and can therefore be compared across all voxels and conditions. The mean activity map was calculated for each frequency presented to crow 1 (Fig. 4B). The spatial distribution of active voxels changed with each of the six stimulus frequencies: low-frequency stimuli elicited a response in more dorso-posterior areas, while higher-frequency stimuli elicited a response in more ventro-anterior areas of the field L complex (Fig. 4B).

We next calculated a frequency map from these activity patterns in crow 1 (Fig. 4C). To do this, we superimposed activity for each sound frequency, using different colors to represent the individual frequencies. In cases where a voxel was active across multiple frequencies, we calculated the weighted average frequency resulting in a detailed frequency map showing clear tonotopy, with a systematically ordered spatial representation of sound frequencies (Fig. 4C). Only a cluster of voxels near the middle showed no positively-correlated activity, separating the otherwise coherently connected topographic map in the field L complex. To further visualize the topographic mapping, we plotted the average activity for each frequency along the dorsoventral axis of the target region (Fig. 4D). These iso-intensity curves show that the location of maximum activity differs between frequencies. Example voxels (n = 6: Fig. 4E) were selected for having the highest change in CBV for each respective pure tone frequency. We performed a one-way ANOVA to determine if the activity of these selected voxels differed between the presented frequencies. For all example voxels, except for the rightmost voxel, the activity elicited by the different frequencies differed significantly (one-way ANOVA, p < 0.001). For the rightmost voxel, the activity did not differ significantly between frequencies (one-way ANOVA, p = 0.1553-generally, the responses elicited by the 8000 Hz condition were lower than for the other lower frequencies).

This procedure and analyses were repeated using six band-pass filtered noise bursts that systematically increased in frequency from 150–350 Hz to 4800–11200 Hz. The tonotopy observed for pure tones could similarly be retraced with band-pass filtered noise (**Fig. S1**). There was a dorso-ventral organization of increasing frequencies.

### Comparison of anesthetized and awake states

Dramatic differences in CBV responses to pure tones and band-pass noise bursts were observed when the crows were awake compared to when they were deeply anesthetized (neurophysiological states). Figure 5A shows the weighted frequency map calculated when crow 1 was deeply anesthetized and Figure 5B is when it was more awake. To compare the number of active voxels between stimulus types, neurophysiological states, and individual birds, we considered a voxel to be active if its CBV activity correlated significantly with the stimulus vector.

**Figure 5:**
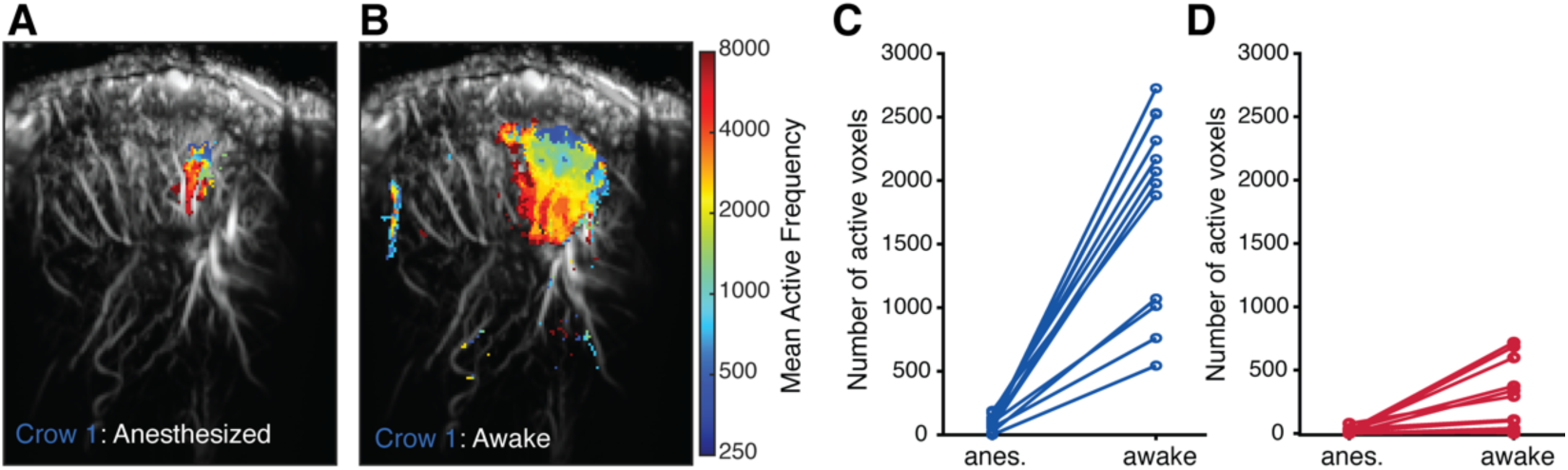
Comparison of auditory activation during neurophysiological states. **A)** Tonotopic frequency map for crow 1 under anesthesia. **B)** Tonotopic frequency map for crow 1 in the awake state. **C)** Number of active voxels in the two different neurophysiological states for crow 1. Each point represents one combination (n = 12 per state) of stimulus type (pure tones, bandpass noise) and stimulus condition (6 frequencies). Lines connect the pairs of the same stimulus condition in the two different neurophysiological states. **D)** Number of active voxels in each neurophysiological state for crow 2.

To investigate the influence of neurophysiological state on the number of active voxels, we counted the number of voxels for each condition (Frequencies 250-8000) and stimulus types (pure tones and band-pass filtered noise) for each bird. This resulted in 12 values for both the anesthetized state and the awake state. For crow 1, an average of 1801.6 ± 217 voxels were active in the awake state, which was significantly more than the 92.2 ± 16.1 active voxels in the anesthetized state (paired t-test, t(11) = 8.2, p < 0.001, Fig. 5C). In crow 2, although there were fewer active voxels in total, the difference between the neurophysiological states was also significant (paired t-test, t(11)= 3.28, p = 0.007, Fig. 5D). In the awake state, we found on average 273.6 ± 78.6 voxels while we found only 19.5 ± 6.6 active voxels in the anesthetized state. Anesthesia clearly led to a reduced overall activity in both birds.

Next, we used MVPA to gain more insight into the stability of the frequency mapping. Since MVPA allows us to use the whole image without preselecting active voxels, it provides a more agnostic analysis approach by utilizing more subtle changes in activity patterns to classify conditions. This makes it particularly useful for datasets with less overall activity. Additionally, since we observed tonotopic mapping in both the anesthetized and awake states, one key question we aimed to resolve was whether this tonotopic pattern was consistent across neurophysiological states. The confusion matrices (Fig. 6A**-D**) show the accuracy with which the classifier predicted conditions for crow 1. Given that there are six frequencies, the chance level is 16.7%.

**Figure 6:**
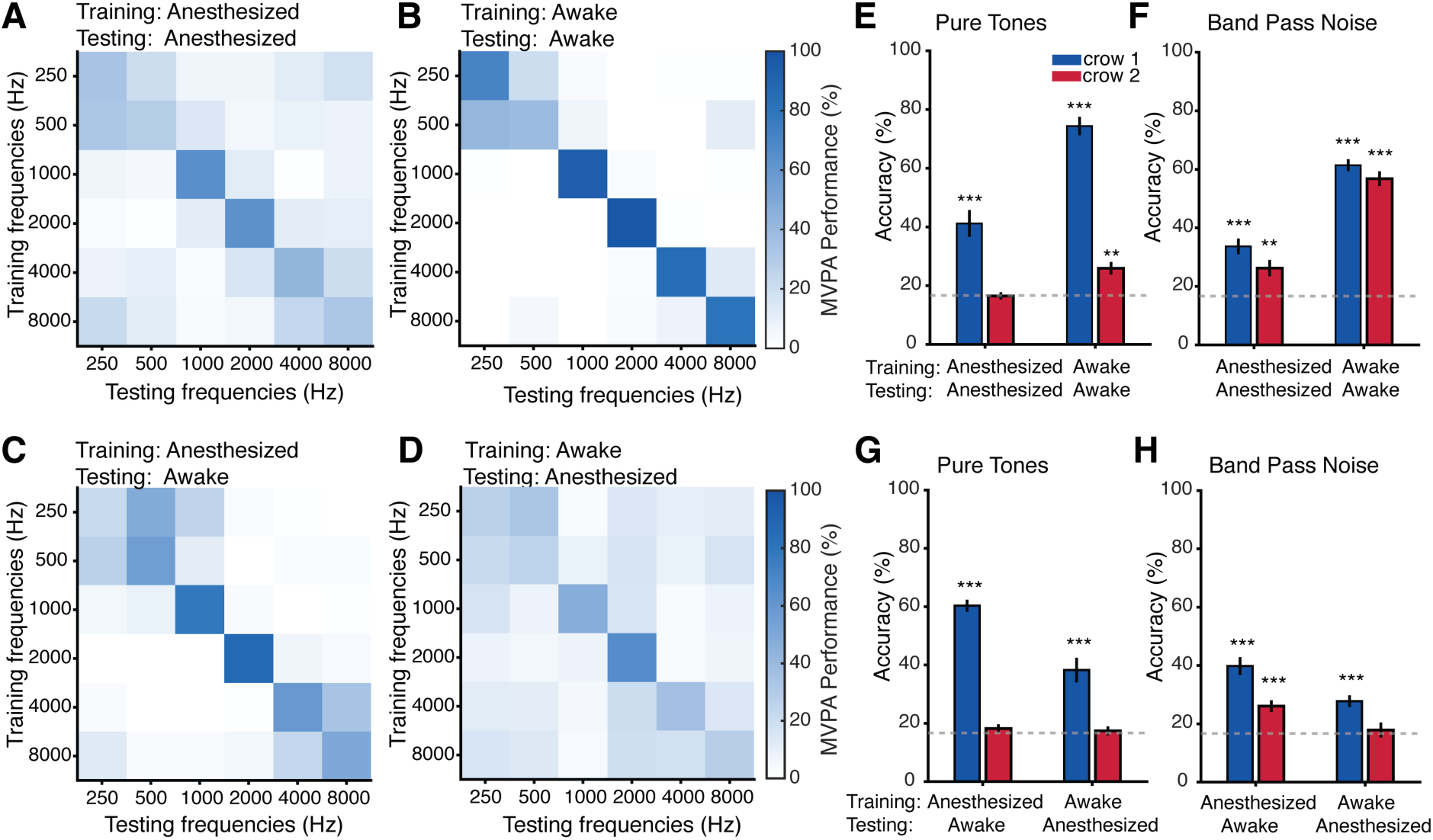
Patterns of voxel activity reveal frequency tuning within and across neurophysiological states. **A)** Confusion matrix displaying the accuracy of an MVPA classifier when predicting the frequency of pure tones played to the anesthetized crow 1. **B)** Confusion matrix for the awake crow 1. **C)** Across-neurophysiological state confusion matrix of the MVPA classifier trained on dataset from the anesthetized crow 1 and tested on data from the awake crow 1. **D)** Across-neurophysiological state confusion matrix of the MVPA classifier trained on data from the awake crow 1 and tested with data from the anesthetized crow 1. **E/F**) Overall accuracy of within-neurophysiological state classifiers for pure tone (**E**) and band-pass noise stimuli **(F**). The grey line indicates chance level for the classifiers given 6 different sound frequencies. Color indicates crow identity (Blue: crow 1, Red: crow 2). **G/H)** Overall accuracy of across-neurophysiological state classifiers for pure tone (**G**) and band-pass noise stimuli (**H**). Error bars indicate the standard error of the mean (SEM). * = p<.05, ** = p <.01, *** p <.001.

Overall, the classifier demonstrated above chance accuracy in predicting the pure tone frequencies for crow 1 (one-sample t-test, p < 0.001) (Fig. 6E, F). It achieved an accuracy of 41.2 ± 4.5% when trained and tested on the anesthetized data (Fig. 6A). Accuracy varied depending on the neurophysiological state, with the highest accuracy observed for the awake-awake condition at 74.4 ± 3.1% (Fig. 6B). Overall, the classifier was able to predict conditions significantly above chance when tested within the same neurophysiological state (Fig. 6E, F). The only exception was the classifier trained and tested on the dataset for pure tone stimulation under anesthesia in crow 2 (t(11) =-0.129, p = 0.9).

The classifiers also generally performed well in predicting sound frequencies when trained across neurophysiological states for crow 1 (Fig. 6G, H) (one-sample t-test, p < 0.001). For the classifier trained on the anesthetized dataset and tested on the awake dataset, accuracy was 58.7 ± 3% (Fig. 6C), while the accuracy dropped to 38.3 ± 4.2% for the classifier trained on the awake dataset and tested on the anesthetized dataset (Fig. 6D). In crow 2, fewer active voxels were observed, and the changes in CBV relative to baseline were less pronounced. The effect of this reduced activity compared to crow 1 is also reflected in the classification accuracy for crow 2. For crow 2, the only across-neurophysiological state classifier that could predict conditions above chance was the one trained on the band-pass noise dataset under anesthesia and tested on the awake dataset (Fig. 6H) (t(11) = 4.723, p < 0.001).

### Effect of stimulus length on CBV changes

Stimulus durations in cognitive tasks are typically shorter than our previously applied 5 second stimuli. We therefore explored CBV responses to three different stimulus durations (5, 1, and 0.5 s) of the same frequency stimulus (1000 Hz pure tone).

We first examined significant correlation maps with a threshold of p < 0.01, FDR corrected (Figure 7A-C) for stimuli lengths of 0.5s, 1s, and 5s, respectively. The correlation maps were robust and similar between the shorter stimuli lengths (0.5s and 1s) compared to the longer stimulus length of 5 seconds. We selected 5 voxels (colored arrowheads in Fig. 7A**-C**) and plotted their CBV traces (Figure 7D-H). CBV responses could be detected for short 0.5 and 1 second stimulations; the magnitude and length of elicited responses were similar between the shorter stimulation lengths. For the two shorter stimulus lengths, CBV increased after stimulation onset to a peak that shortly decayed. Interestingly, the responses differed with the longer 5s stimulation duration. In **Figure 7D, G and H**, the example voxels (orange, magenta and purple arrowheads) had sustained activity throughout the entire stimulation period and past the offset of the sound. In Figure 7E, the example voxel (yellow arrowhead) had a short increase in CBV before a larger, longer negative deflection. In Figure 7F, the example voxel (red arrowhead) showed a transient positive response at the onset and the offset of the stimulus. This diversity of responses suggests that fUSi could be used to examine putative functional subfields within the field L complex that show different CBV changes with time.

**Figure 7:**
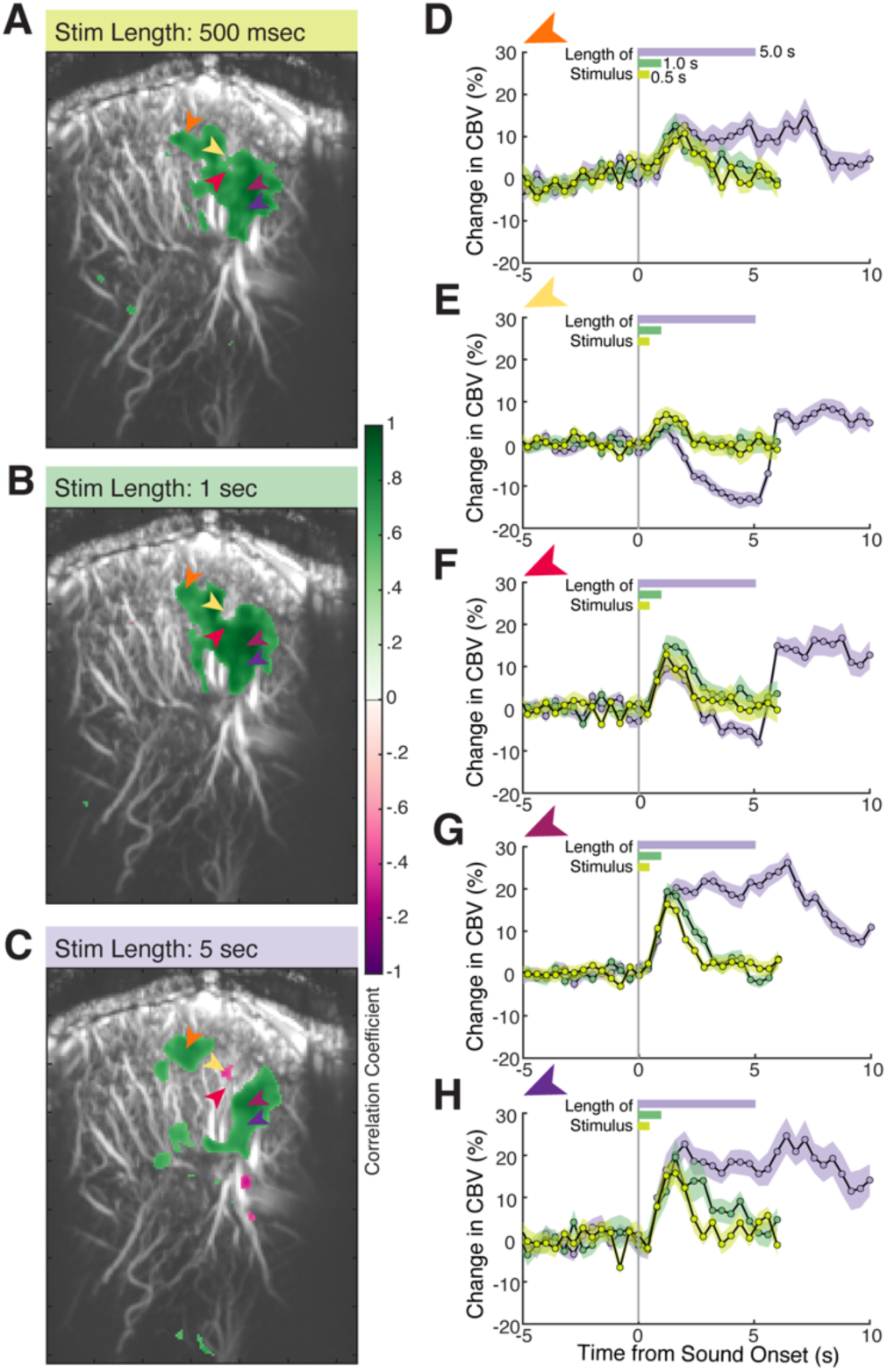
Auditory responses in the field L complex are scaled by length of stimuli presented. **A)** Correlation map of active voxels to a 0.5s stimulus (Pure tone: 1000Hz). Positive correlations are in green and negative correlations in pink. The five colored arrowheads indicate voxels that have associated CBV traces plotted in (**D-H**). **B)** Correlation map to a 1s stimulus. **C)** Correlation map to a 5s stimulus. **D)** The activity of the corresponding voxel (orange arrowhead) for playback of a 500 ms (light green), 1 s (dark green), and 5 s (purple) 1000Hz pure tone. On the x-axis, zero is sound onset. Note the differing length of CBV traces is due to the addition of 5 s of scan time post stimulus offset. **E-H)** Other example voxels plotted similarly to (**D**) that span the correlation map from top left to bottom right.

To examine these dynamics, we superimpose the CBV value for each voxel that passed a threshold of 5% change from baseline for each fUS image (0.4s duration), starting from 0.8s before stimulus onset to 1.4s after the end of the 5s sound (Figure 8). Different patterns of activity emerge during the onset, maintenance, and offset of the 5s sound stimulus. For the first few frames after stimulation onset, there is a positive increase throughout the field L complex. After about 2 seconds, the region appears to divide with two flanking robust sustained positive responses and a middle region that experiences a negative deflection. After the offset of the stimulus, this negative deflect reduces and the divisions appear to merge again into a cluster of positive activity. This visualization suggests that different response dynamics could indicate finer spatial organization within the field L complex.

**Figure 8:**
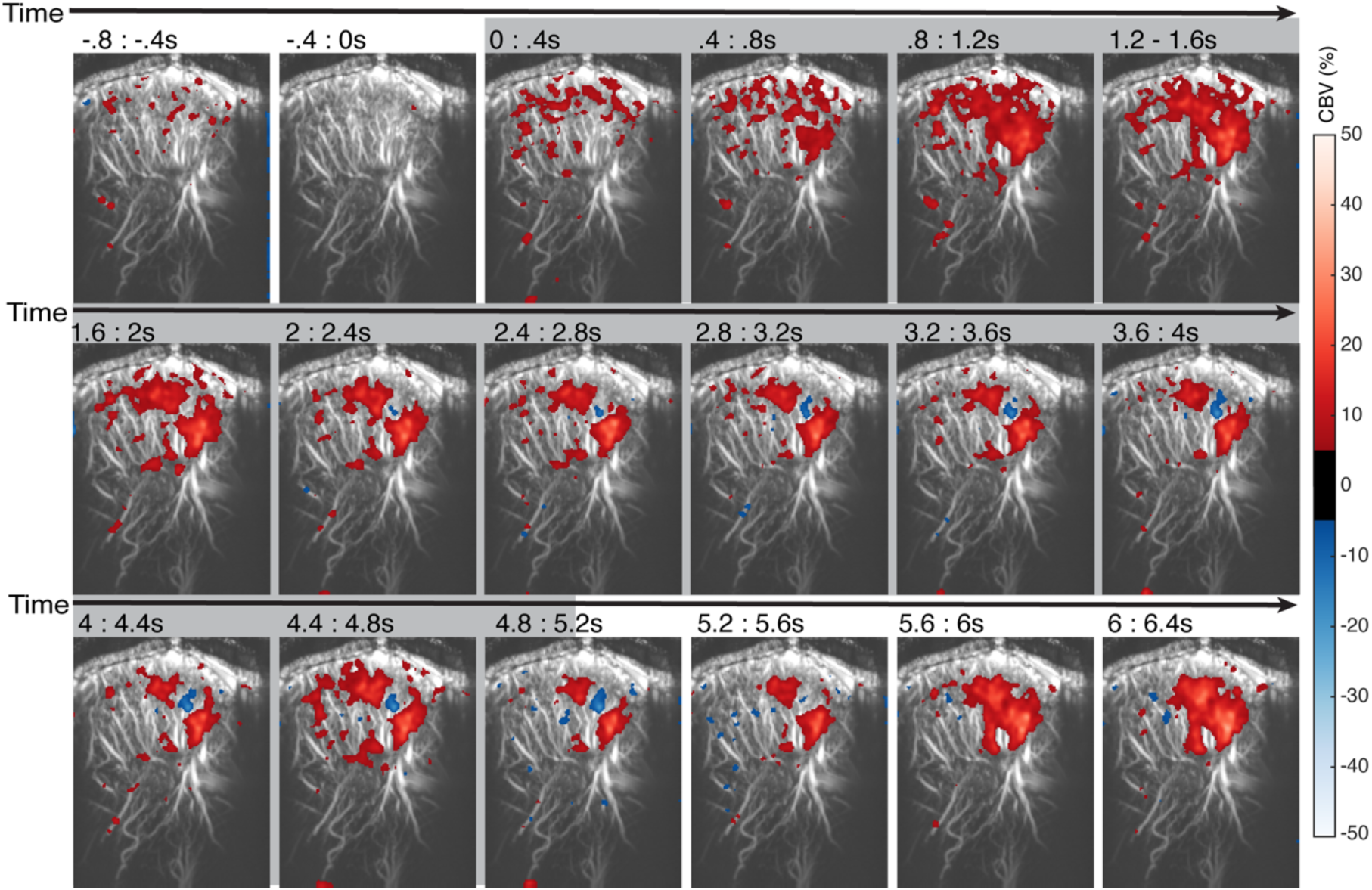
Dynamics of auditory responses in the field L complex to a 5 second stimulus. Each fUS image spanning a stack of compound images over 0.4s is plotted from 2 images before the onset of the sound stimulus to 3 images after the offset of the sound stimulus for crow 1. The percent change in CBV (thresholded by 5%) is plotted for each image. Different patterns of activity emerge during the sound stimulus (as marked by the gray background) and afterwards as images progress through time from left to right along each row. Positive CBV changes are colored in red and negative CBV changes are colored in blue.

## Discussion

We applied functional ultrasound imaging (fUSi) to the crow brain. We could generate detailed 3D maps of the crows brain vasculature and then map out the location of the field L complex within the left hemisphere with white noise stimulation. Investigating field L complex responses to naturalistic, complex sound stimuli revealed a spatial organization informative of the stimuli’s frequency content, allowing reliable classification of three different bird vocalizations with MVPA. A fine-scale topographic analysis showed robust tonotopic mapping along the dorsoventral axis of the field L complex, with lower frequencies represented dorsally and higher frequencies ventrally. Due to the high signal to noise ratio, voxel-specific analysis could demonstrate sharp frequency tunings consistent with this tonotopy. Notably, the tonotopic organization persisted across neurophysiological states, and MVPA revealed the robustness of this mapping across states. We were further interested in the temporal responses to sensory stimuli of different lengths. We found that short stimuli robustly elicited a peak whereas longer stimuli yielded more complex response patterns. When we visualized the activity in a time resolved manner, different spatial regions appeared to have different response patterns, suggesting smaller subregions within the field L complex.

This study represents the first application of fUSi to investigate the songbird brain. Previously, Rau et al. (2018) demonstrated the feasibility of fUSi in birds, localizing the pigeon visual wulst and entopallium via visual stimuli and the field L complex via auditory stimuli. Here, we provide a detailed investigation into the dynamics of the field L complex’s response properties using diverse auditory stimuli, including naturalistic and frequency-controlled sounds, comparing across anesthetized and awake states.

### Methodological considerations

Our imaging procedures left the inner bone layer intact while achieving high-quality imaging, allowing us to reconstruct detailed tonotopic maps with as few as six to twelve scans. This efficiency allowed us to map the response to auditory stimuli in one slice in as little time as 15-30 minutes, enabling us to examine the effect of anesthesia on the representation of frequencies all in one session. The amplitude of CBV signals obtained from crow fUSi were around 30% and thus large compared to typical BOLD responses obtained from fMRI (Boido et al., 2019). Similarly, auditory responses have been reported to be ∼30% in the ferret (Bimbard et al., 2018), and ∼20-30% in marmosets (Takahashi et al., bioRxiv) compared to typical auditory cortex BOLD responses of ∼5%. FMRI in zebra finches has shown that it is possible to locate the position of the field L complex in the birds’ brains, but at a lower spatial resolution in a smaller bird (Arya et al., 2023). However, fUS images are 2D. 3D imaging (Gesnik et al., 2017, Rau et al., 2018, Bimbard et al., 2018, Rabut et al., 2019, Brunner et al., 2020, Bertolo et al., 2021) currently requires the use of heavier motors or larger probes, precluding a head-free protocol in the crow. More lightweight volumetric probes are actively being developed.

Currently, fUSi allows a wide field of view, high spatial resolution and a temporal resolution fast enough to image brain dynamics underlying complex behaviors and task-relevant intervals. This has led to its application to diverse species: mice (Macé et al., 2011,Aydin et al., 2020, Bertolo et al., 2021, Boido et al., 2019, Brunner et al., 2020, Edelman et al., 2021, Lambert et al., bioRxiv), rats (El Hady et al., 2024, Sieu et al., 2015, Tiran et al., 2017, Urban et al., 2015, Rabut et al., 2019), macaques (Claron et al., 2023, Dizeux et al., 2019, Norman et al., 2021, Griggs et al., 2024), marmosets (Takahashi et al., bioRxiv, Zhang et al., 2022), ferrets (Hu et al., 2023, Bimbard et al., 2018), pigeons (Rau et al., 2018), newborn and adult humans (Demene et al., 2017, Imbault et al., 2017, Rabut et al., 2024). Critically, fUSi can be performed in awake and even freely-moving animals to study more complex behaviors (Brunner et al., 2020, Urban et al., 2015, Sieu et al., 2015, Tiran et al., 2017, Takahashi et al., bioRxiv, El Hady et al, 2024). This makes it a highly attractive technique to study the complex cognitive and communicative behaviors of corvids.

### CBV changes in crows reflect neural activity in the field L complex

The functional neuroanatomy and physiology of the auditory telencephalon, specifically the field L complex, in corvids had previously been unexplored. The field L complex is the primary auditory area in the avian pallium, functionally analogous to the primary auditory cortex in mammals (Elliott & Theunissen, 2011). This region plays a crucial role in processing external sounds, including vocalizations and songs, as well as self-generated vocalizations.

In our study, we observed sound-induced changes in CBV in the posterior telencephalon of crows. Notably, the anatomical coordinates of these CBV changes align with the anatomical coordinates of the field L complex, as reported in previous histological studies (Kersten et al., 2021; 2022; 2024). Additionally, the location of CBV changes observed in our study corresponds to regions where neuronal activity in response to both simple and complex sounds has been recorded in other songbirds, such as zebra finches (Gehr et al., 1999; Calabrese & Woolley, 2015) and starlings (Müller & Leppelsack, 1985; Rübsamen & Dörrscheidt, 1986). This anatomical and physiological correspondence suggests that the observed CBV changes in crows support the functional similarity of the field L region across different avian species.

The field L complex in birds, while varying in specific subdivisions and terminology across species, is generally agreed to be organized into three distinct, sandwiched layers: L1, L2, and L3 (Bonke et al., 1979; Fortune & Margoliash, 1992; Vates et al., 1996). L2 serves as the primary recipient of thalamic input from the auditory nucleus ovoidalis (Karten, 1968; Wang et al., 2010). From L2, auditory information is processed and relayed to the other two subregions, L1 and L3. L1 and L3 serve as the output layers of the field L complex, sending information to regions such as the nidopallium caudo-medial (NCM) and the caudal mesopallium (CM), which are involved in song memory, recognition, and learning (Bailey et al., 2002; Thompson & Gentner, 2010). In oscine birds, including crows, L1 and L3 also project to the dorsal nidopallium near the song nucleus HVC, particularly to the HVC shelf, as well as to the nidopallium caudolaterale (NCL), an executive center involved in higher-level processing (Kersten et al., 2024; Moll et al., 2024; Vates et al., 1996). These projections highlight the integrative role of the field L complex in auditory processing and song-related behaviors.

### Tonotopic arrangement in the corvid field L complex

A notable finding in this imaging study is the systematic regional variation in CBV based on sound frequency. Low-frequency sounds are represented in the posterior-dorsal region, while progressively higher frequencies are mapped towards the anterior-ventral region. This tonotopic organization in the field L complex resembling the organization of the mammalian auditory cortex, where neurons are systematically arranged according to their characteristic frequency responses to form a tonotopic map (Elliott & Theunissen, 2011).

In non-corvid songbirds, neurons in the field L complex show overlapping frequency tuning curves, which together create an integrated tonotopic representation. Iso-frequency contours in the field L complex run perpendicular to the L2 subfield and span all three subfield layers (L1, L2, and L3), as demonstrated by earlier electrophysiological studies (Müller & Leppelsack, 1985). In species like European starlings and zebra finches, each of these subfield layers forms a tonotopic map of the basilar papilla, organizing sound frequencies from low to high (Gehr et al., 1999; Capsius & Leppelsack, 1999).Specifically, the cochleotopic gradient maps lower frequencies to the anterio-dorsal region and higher frequencies to the postero-ventral region of the field L complex (Müller & Leppelsack, 1985; Rübsamen & Dörrscheidt, 1986).

The findings of the present study, derived using functional ultrasound (fUS) imaging, are consistent with electrophysiological results previously observed in non-corvid songbirds, suggesting a similar tonotopic organization in the corvid field L complex. Additionally, fUSi demonstrated its capability to detect CBV changes with spatial and temporal resolution of 100µm and several 100 ms, highlighting its utility for detailed investigations of the auditory system. Notably, fUSi has also been used to produce stable, high-resolution tonotopic maps in the auditory pathways of awake ferrets (Bimbard et al., 2018), further underscoring its versatility and precision.

### Processing hierarchy within the corvid field L complex

The comparison of CBV changes under different neurophysiological states, along with the patterns observed during prolonged sound stimulation, suggests that fUS can be used to delineate between putative subregions of the field L complex (Elliott & Theunissen, 2011). Under anesthesia, CBV responses were confined to the central region of field L, which would correspond to the thalamo-recipient input region, L2. This region still maintained a topographic layout, suggesting that only L2 responded to sounds, while the flanking postsynaptic and higher-order auditory regions, L1 and L3, were largely inactive.

Interestingly, this central part of the field L complex (interpreted as L2) exhibited a brief phasic excitation immediately after stimulus onset, followed by a pronounced suppression below baseline during prolonged sound stimulation. In contrast, the surrounding parts (interpreted as L1 and L3) showed a sustained increase in CBV throughout the stimulation period. This spatially dissociated pattern of CBV changes resulted in a marked dissociation between the anterior (proposed L1) and posterior (proposed L3) regions, with a negative or brief onset and offset CBV changes in the center (L2).

This proposed delineation is consistent with the electrophysiological properties of neurons in these regions. Neurons in L2, which receive direct thalamic input, exhibit the shortest response latencies. In contrast, neurons in L1 and L3, along with those in the CM and NCM areas, show progressively longer latencies (Calabrese & Woolley, 2016). This latency pattern supports the notion of a feed-forward processing hierarchy, with L2 as the initial processing center, followed by the more integrative roles of L1 and L3 (Wang et al., 2010; Calabrese & Woolley, 2016). The observed processing hierarchy in the field L complex mirrors the principles of information processing seen in the canonical cortical microcircuit, which was once thought to be exclusive to mammals (Calabrese & Woolley, 2016).

### Influence of anesthesia on auditory-induced CBV

We observed a significant reduction in stimulus-induced CBV changes in the auditory telencephalon of crows under ketamine/xylazine anesthesia compared to the awake state. This aligns with findings from animal fMRI studies, which have shown that awake animals exhibit stronger BOLD responses compared to those under anesthesia (Desai et al., 2011; Dinh et al., 2021). Studies on auditory processing across different brain levels and under various anesthetics consistently demonstrate that neuronal responses are significantly altered during anesthesia. Evidence from human imaging studies (Heinke and Koelschb, 2005) and animal electrophysiological recordings (Gaese and Ostwald, 2001; Syka et al., 2005) highlights these changes. In songbirds, research has shown that both spontaneous and stimulus-induced neuronal activity in the field L complex are significantly reduced under anesthesia compared to the awake state (Capsius and Leppelsack, 1996; Schmidt and Konishi, 1998; Karino et al., 2016). This suppression of activity is a common finding in anesthetized animals, indicating that anesthesia broadly diminishes auditory processing.

In our study, we used ketamine combined with xylazine as the primary anesthetic agents. Ketamine functions primarily as an N-methyl-d-aspartate receptor (NMDAR) antagonist, blocking excitatory glutamatergic synaptic transmission (Zanos et al., 2018). The most straightforward interpretation of our findings is that the substantial reduction in stimulus-induced CBV changes observed in crows under anesthesia reflects reduced neuronal activity in the field L complex. However, we cannot entirely rule out the possibility that the observed CBV reduction is influenced by the direct effects of anesthesia on cerebral blood flow (CBF) via mechanisms affecting the cerebral vasculature. Regardless of the exact underlying mechanism, it is clear that anesthesia leads to a significant underestimation of functional auditory activation.

The classifier’s superior performance in across-neurophysiological state classification highlights the preserved stability of tonotopic mapping under anesthesia. Although anesthesia suppresses overall neuronal activity, the underlying tonotopic organization remains intact. This allows classifiers trained on awake-state data to predict anesthetized conditions with moderate accuracy, despite reduced activity levels. In contrast, classifiers trained on anesthetized data achieve higher accuracy when tested on awake data due to the increased activity in the latter state. These findings demonstrate that while anesthesia reduces neural activation, it does not disrupt the fundamental organizational features of the auditory telencephalon.

### Effects of stimulus duration on CBV

We analyzed a critical parameter for behavioral and physiological studies-the effects of stimulus duration-on the recorded CBV. Our data demonstrate that CBV changes can be reliably detected even with shorter stimuli of 0.5s and 1s lengths, and these changes are consistently observed at the single-voxel level. This capability is particularly advantageous for studies involving behaviorally trained animals, where short stimuli in the range of 500 ms are commonly employed. Following stimulus onset, the CBV signal shows a transient increase that quickly decays. In contrast, longer stimuli (e.g., 5s) produces more variable responses, including sustained activity, onset-specific responses, and even occasional negative deflections. It is worth mentioning that the widely used methods to identify significantly active voxels by correlating CBV signals with a convolved stimulus vector may miss such complex stimulus evoked activity that does not follow the shape of the stimulus vector in a simple manner.

We found that amplitudes of initial responses were similar across stimuli durations, indicating robust signal-to-noise ratios. However, differences emerged after the first few seconds when longer stimuli of 5 seconds were presented. Similarly, studies in macaque monkeys have reported comparable response profiles in V1 activity for 0.5s and 1.0s visual stimuli (Blaize et al., 2019). In behavioral tasks, shorter stimuli are often sufficient to capture relevant activity (Norman et al., 2021, Griggs et al., 2023, El Hady et al., 2024), even allowing online decoding of eye movement direction during a memory-guided saccade tasks (Norman et al., 2021, Griggs et al., 2023).

We observed a significant negative response during presentation of a 5s long stimulus that was absent in 0.5 and 1.0-second stimuli scans. Consistent negative responses have been reported in other investigations, including olfactory (Boido et al., 2019) and auditory (Takahashi et al., bioRxiv) studies. Such negative responses could result from factors like artery constriction, steal effects, or drainage. In fUSi recordings of pigeon field L, a negative CBV change was localized to an arteriole supplying the deeper capillary structure of the brain, near where a larger positively changing regions was observed (Rau et al., 2018); in this study, the auditory stimuli used were hawk calls and frequency-matched noise, each played for 15 seconds.

## Conclusion

In conclusion, our study establishes functional ultrasound imaging (fUSi) as a powerful tool for studying auditory processing in crows. fUSi enables high-resolution mapping of CBV changes in the field L complex, revealing tonotopic organization and subregional responses to diverse auditory stimuli. Its sensitivity across stimulus durations and neurophysiological states underscores its versatility for investigating neural dynamics. By bridging neural activity, vascular responses, and behavior, fUSi offers valuable insights into avian functional systems and facilitates comparative studies across species.

## Acknowledgments

We thank Philipp Nieder for 3D-printing implants, Alexander Song for help with data analysis, Ylva Kersten for useful discussions on crow neuroanatomy, Haleh Soleimanzad and Sara Romanzi for fUS training sessions.

## Author Contributions

Conceptualization: D.A.L., E.S., A.N.

Methodology: D.A.L., E.S., A.N.

Investigation: D.A.L., E.S., A.N.

Visualization: D.A.L., E.S.

Supervision: A.N.

Writing—original draft: D.A.L., E.S., A.N.

Writing—review & editing: D.A.L., E.S., A.N.

## Methods

### Animals

Data was acquired for two female carrion crows (*Corvus corone*) from the institute’s facility. Crows were housed in small social groups in large indoor and outdoor aviaries (for additional details see Hoffmann et al., 2011). Crows were provided food and water ad libitum. The night before experimental days, crows were fasted for anesthetic purposes. Crow 1 was 1 year old and participated in two sessions separated by 12 days. Crow 2 was 8 years old and participated in a single session. All procedures were conducted in accordance with German and European law and approved by the local authority, the Regierungspräsidium Tübingen.

### Functional Ultrasound Imaging

Functional ultrasound data were acquired using the Iconeus One system (Iconeus, Paris, France), specifically designed for animal studies, as described previously (Bertolo et al., 2021). A linear ultrasonic probe (IcoPrime-15 MHz, Iconeus, Paris, France) was connected to the ultrafast ultrasound scanner (Iconeus One, 128 channels). The probe, consisting of a linear array of 128 piezoelectric elements with a 15 MHz center frequency and a 0.1 mm pitch, provided a broad field of view (14 mm width, up to 20 mm depth, and 400 μm plane thickness) with an in-plane spatial resolution of 100 μm × 100 μm.

2D fUS imaging was conducted at a frame rates of 2.5 Hz. Power Doppler (PD) images were generated from 200 compounded frames acquired at a rate of 500 Hz to form an PD image every 400msec. The ultrasound sequence involved transmitting 11 tilted plane waves (ranging from −10° to 10° in 2° increments) at a pulse repetition frequency (PRF) of 5.5 kHz. Each block was processed using Singular Value Decomposition (SVD) clutter filtering to separate tissue signals from blood signals, resulting in the formation of a PD image (Demené et al., 2015). The custom imaging sequence and real-time Doppler reconstruction were executed using dedicated acquisition software (IcoScan, Iconeus, Paris, France).

### Surgical Procedures

All surgeries were performed while the animals were under general anesthesia. Crows were anesthetized with a ketamine/xylazine mixture (50 mg ketamine, 5 mg xylazine /kg initially and supplemented by 17 mg ketamine, 1.7 mg xylazine /kg i.m. per hour during the ongoing surgery). After the surgery, the crows received analgesics (Butorphanol (Morphasol®), 1 mg/kg i.m.). The head was placed in the stereotaxic holder that was customized for crows with the anterior fixation point (i.e., beak bar position) 45° below the horizontal axis of the instrument. The skull overlying the brain region of interest was exposed by making an incision along the top of the head through the skin. In birds, the skull is composed of three main layers: the outer cortical bone layer, which is a compact outermost layer; the trabecular bone (spongy layer) beneath the cortical bone; and the compact inner bone layer surrounding the brain. A partial craniotomy (dimensions 18 mm x 18 mm) was performed over the left posterior telencephalon, extending across the midline a few millimeters into the right hemisphere. In a partial craniotomy, the outer cortical bone and trabecular bone are removed, while the inner bone layer is left intact. This inner bone layer is thin, allowing for imaging through it without the probe making direct contact with the brain. The same type of partial craniotomy has proven successful for fUSi in pigeons (Rau et al., 2018). The area of interest in the center of the partial craniotomy was the field L complex, the telencephalic auditory area in birds, which in crows has been localized to 2-3 mm from the midline and 5 mm anterior to the bifurcation of the sagittal sinus (Kersten et al., 2022).

A custom 3D printed basin was attached to the intact bone surrounding the partial craniotomy site with dental cement. The basin was designed with ridges that allow for the attachment of a fUS probe holder during recording sessions or a flat cover for after recording sessions to protect the site. During recording, ultrasound transmission gel (Aquasonic, Parker Laboratories, Inc.) was used to fill the basin in which the probe was positioned and secured.

### 3D Angiogram

3D angiogram recordings of the brain’s vasculature were carried out using the integrated “3D angiogram” function of IcoScan with a linear motor stage (SLC-1740, SmarAct GmbH). The motor was placed over the craniotomy and attached to a probe holder. The motor-coupled probe could then be moved over the craniotomy in either the mediolateral axis or the anterior-posterior axis to acquire sagittal or coronal scans respectively. The 3D scan was carried out in step sizes of 0.2 mm.

### Localization of field L recording slice

For imaging auditory responses, we positioned the probe in the sagittal orientation, recording a 2D slice with a field of view of 14mm x 19mm. To determine the best slice to elicit auditory responses, we targeted the region 2-3 mm from the midline. In the surgical suite, a broadband sound was played to the bird for each slice, to ensure the correct probe placement before transferring the bird to the recording setup. With IcoStudio, the correlation maps between the stimulus vector and the power Doppler signals were computed. We noted down which slices showed the highest correlation values with sound presentation and fixed the probe to the slice position with the highest and largest activity. We then transfered the bird to the acoustic setup with the probe fastened to the basin. Upon arriving in the recording chamber we immediately checked the position of the probe again by repeating the previous stimulation pattern and continued with planned auditory sequences.

### Sound playback and recording apparatus

The experiment was conducted in a ventilated double-walled sound-proof chamber (IAC Acoustics, Niederkrüchten, Germany). A linear broadband speaker (Visaton) played auditory stimuli at an average sound pressure level of 65 dB. The bird was placed in a wire enclosure, approximately 50 cm from the speaker. The chamber was equipped with a camera using infrared LEDs to monitor the bird’s behavior throughout the experiment. No visible light was used in the chamber.

The Iconeus One system was arranged outside the chamber to allow the probe to be attached to the bird’s head via the probe holder, with sufficient flexibility to accommodate head movements. A split BNC cable split the audio output to be played on the speaker in the recording chamber and to the Iconeus One system. A pure tone pulse at the beginning of the sound file, not audible on the speaker in the recording chamber, triggered the start of acquisition. This ensured that recordings could be aligned to the stimulation.

### Stimulus sequences

Ultrasound recordings were performed in two different neurophysiological states. In the anesthetized state, the crows were fully sedated with fresh injections of the ketamine/xylazine anesthetic, resulting in loss of consciousness, absence of reflexes, immobility, and muscle relaxation. After several hours, when the effects of the anesthesia wore off, the crows were imaged in the awake state. In this state, the crows maintained a stable posture on their feet, exhibited spontaneous movements such as shifting positions and head movements, kept their eyes open, and responded to stimuli, all while resting calmly in a darkened, soundproof chamber.

In the experiment, we used four distinct stimulus sets to investigate auditory responses. The first set comprised of complex sounds, specifically vocalizations from a crow, pigeon, and canary. Each sequence featured 5 seconds of vocalizations from each bird species, followed by a pause of 15 seconds, with the order of vocalizations pseudorandomized. The calls varied in frequency content.

Then, pure tones and band-pass filtered noise were employed to characterize tonotopic mapping. The second stimulus set consisted of six pure tones, logarithmically spaced from 250 Hz to 8000 Hz, covering the hearing range of carrion crows (Jensen & Klokker, 2006). Each pure tone was presented for 5 seconds, followed by a 15-second silence. The six tones were played in sequence, constituting one acquisition, with a total duration of 135 seconds, including a 15-second baseline period in the beginning of the acquisition. The order of stimuli was pseudorandomized across sequences so that each condition was presented once per scan, aiming for 12 scans for a complete dataset.

The third stimulus set involved band-pass filtered noise, with center frequencies corresponding to the pure tones in the previous set. The frequency bands increased in width with higher frequencies, ranging from 150-300 Hz for the lowest frequency condition to 4800-11200 Hz for the highest frequency condition. The randomization protocol was the same as for the pure tones.

To determine the response to different stimulus duration, we presented a 1 kHz pure tone for 0.5, 1, and 5 seconds. This frequency was chosen based on previous experiments, where it elicited a stronger response and is well-represented in the crow’s auditory system (Jensen & Klokker, 2006).

### Selection of Active Voxels

Scans were converted to.mat files for further processing. Analyses were conducted using MATLAB (Version R2023a, MathWorks Inc., Natick, MA). First, slices in time were corrected with a non rigid motion correction algorithm (NoRMCorre, Pnevmatikakis & Giovannucci, 2017). Individual voxels were spatially smoothed with a Gaussian kernel of 3 voxel widths. Then, for each voxel, the normalized change in cerebral blood volume (CBV) was calculated as a percentage by subtracting the mean baseline value from the trace and dividing the result by the mean of the baseline.

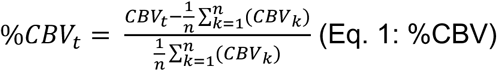

The baseline was defined as the period 5 seconds before stimulus onset up to the onset itself. To identify active voxels, we first created a binary stimulus vector, where for example with a 5 second length stimulus, the 5-second off-phase before stimulus onset was labeled as 0, and the 5-second stimulation phase was labeled as 1. This binary stimulus vector was then convolved with the hemodynamic response function (HRF). The HRF was obtained by fitting an inverse gamma distribution to the data (Eq. 2).

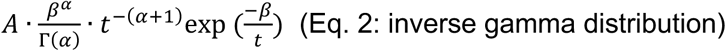

The resulting HRF was then convolved with the binary stimulus vector. The resulting convolved stimulus vector was then used to identify active voxels. To do so, the change in CBV during stimulation was correlated with the convolved stimulus vector using Pearson correlation. To control for multiple comparisons, the Benjamini-Hochberg procedure was applied for False Discovery Rate (FDR) correction, with voxels showing a corrected p-value less than 0.001 classified as active.

### Frequency Maps

We calculated the frequency maps by noting down the frequency which elicited a response for each significant voxel. In cases where a voxel was active for multiple frequencies, we calculated a weighted average from those frequencies. The weighted average was obtained by normalizing the activity each frequency elicited in the specific voxels to the maximum activity. The normalized activity was then multiplied with the respective frequency. Finally, the resulting proportioned frequencies were averaged per voxel.

### Multivoxel Pattern Analysis (MVPA)

We used the Princeton MVPA toolbox (Detre et al, 2006, Norman et al., 2006) to conduct the multi-voxel pattern analysis. The code, originally designed for fMRI datasets, was adapted for fUS-acquisition data files as input and was implemented into a custom script. The MVPA was conducted on the raw data, i.e. without normalization to baseline, spatiotemporal smoothing or preselection of voxels. Only timepoints that fall into the shifted stimulation period were fed into the MVPA. The input data were z-scored independently for each run. In anticipation of a leave-on-out cross validation scheme, the data was sorted into folds, with each fold corresponding to one trial sequence. In each iteration of the cross-validation, one fold was used as testing data and n-1 folds were used as training data. In each iteration a feature selection using an ANOVA with a p < 0.05 threshold was performed. The ANOVA selected voxels which were significantly different between the conditions. Non-significantly different voxels were excluded from the analysis. This feature selection was performed only on the training data and individually for each iteration to ensure that no bias is introduced by manually selecting active voxels. In each iteration a backpropagation classifier with 10 hidden layers was trained on n-1 scans as training data and 1 scan as testing data. We report the mean accuracy and standard error of the classifiers.

---

**Figure S1:**
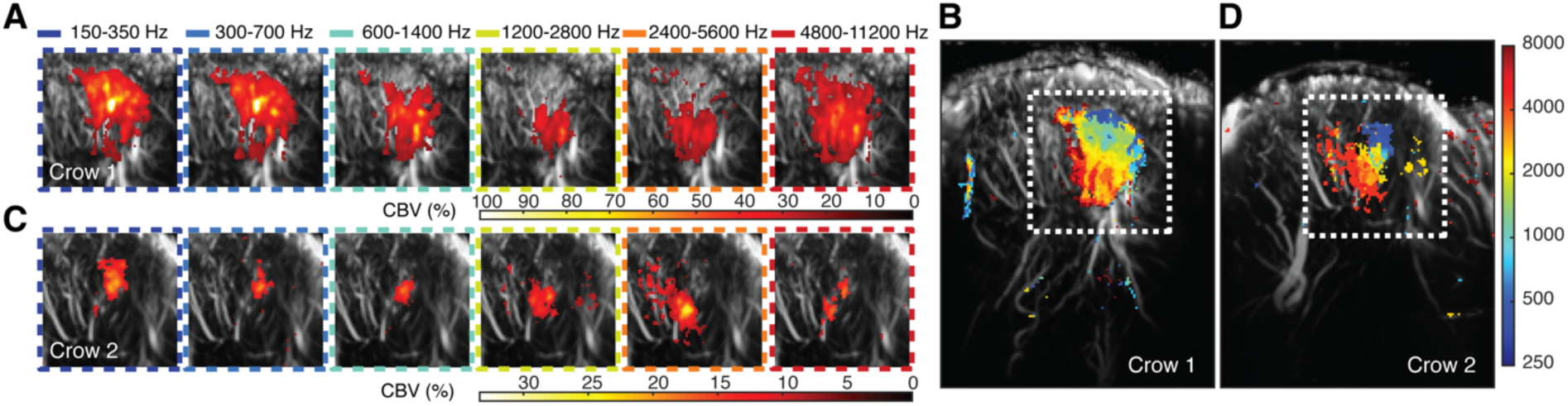
Topographic arrangement of band-pass noise representations in awake crows. **A)** Average activation maps in crow 1 (graphically zoomed in on the field L complex; see dotted white square in **B** show increasing band-pass noise frequencies arranged from left to right. The color of the frames indicates the sound frequency. **B)** Tonotopic frequency map in crow 1. Activation to each band-pass noise sound was superimposed and a weighted averaged was computed, with cool to warm colors representing increasing sound frequencies. **C)** The average activation maps to increasing band-pass noise frequencies for crow 2. **D)** Tonotopic frequency map in crow 2.

